# Distinct features of nucleolus-associated domains in mouse embryonic stem cells

**DOI:** 10.1101/740480

**Authors:** Aizhan Bizhanova, Aimin Yan, Jun Yu, Lihua Julie Zhu, Paul D. Kaufman

**Author notes:** Contact;, University of Massachusetts Medical School, Department of Molecular, Cellular and Cancer Biology, 364 Plantation St. Worcester, MA 01605 USA.

## Abstract

**Background:** Heterochromatin in eukaryotic interphase cells frequently localizes to the nucleolar periphery (nucleolus-associated domains, NADs) and the nuclear lamina (lamina-associated domains, LADs). Gene expression in somatic cell NADs is generally low, but NADs have not been characterized in mammalian stem cells.

**Results:** Here, we generated the first genome-wide map of NADs in mouse embryonic stem cells (mESCs) via deep sequencing of chromatin associated with biochemically-purified nucleoli. As we had observed in mouse embryonic fibroblasts (MEFs), the large Type I subset of NADs overlaps with constitutive LADs and is enriched for features of constitutive heterochromatin, including late replication timing and low gene density and expression levels. Conversely, the Type II NAD subset overlaps with loci that are not lamina-associated, but in mESCs, Type II NADs are much less abundant than in MEFs. mESC NADs are also much less enriched in H3K27me3 modified regions than are NADs in MEFs. Additionally, comparision of MEF and mESC NADs revealed enrichment of developmentally regulated genes in cell type-specific NADs. Together, these data indicate that NADs are a developmentally dynamic component of heterochromatin.

**Conclusions:** These studies implicate association with the nucleolar periphery as a mechanism for developmentally-regulated gene silencing, and will facilitate future studies of NADs during mESC differentiation.

## Introduction

Eukaryotic genomes are broadly subdivided into more accessible, transcriptionally active euchromatin, and less accessible, less active heterochromatin. These functional classifications are accompanied by spatial separation: heterochromatin is mainly found at the nuclear periphery and nucleolar periphery, where they comprise nucleolus-associated domains (NADs) (Németh et al. 2010; van Koningsbruggen et al. 2010) and lamina-associated domains (LADs) (Pickersgill et al. 2006; Guelen et al. 2008; Peric-Hupkes et al. 2010), respectively. Studies in multiple organisms indicate that sequestration of heterochromatin to the nuclear and nucleolar peripheries contributes to gene silencing (Fedoriw et al. 2012b; Zullo et al. 2012; Jakociunas et al. 2013). Therefore, there is great interest in discovering the molecular bases for these localizations. Notably, some trans-acting factors that specifically affect lamina (Zullo et al. 2012; Harr et al. 2015) or nucleolar (Yusufzai et al. 2004; Zhang et al. 2007; Mohammad et al. 2008; Padeken and Heun 2014; Smith et al. 2014; Matheson and Kaufman 2017; Singh et al. 2018) associations have been reported, suggesting that distinct mechanisms contribute at the two locations.

Both NADs and LADs are enriched for silent genes and histone modifications characteristic of constitutive heterochromatin, e.g. H3K9me2 and H3K9me3 (Matheson and Kaufman 2016; van Steensel and Belmont 2017). LADs have been mapped and studied in multiple species and cell types (Pickersgill et al. 2006; Guelen et al. 2008; Peric-Hupkes et al. 2010; Kind et al. 2013; Borsos et al. 2019). In contrast, NADs have been characterized in a few human somatic cell lines (Németh et al. 2010; van Koningsbruggen et al. 2010; Dillinger et al. 2017), in the plant *Arabidopsis thaliana* (Pontvianne et al. 2016), and recently, in mouse embryonic fibroblasts (MEFs) (Vertii et al. 2019). Several experiments indicate that LADs can be redistributed to the nucleolar periphery after passage through mitosis, and vice versa (van Koningsbruggen et al. 2010; Kind et al. 2013). However, the extent of overlap between LADs and NADs is unknown in most organisms and cell types.

Here, we mapped and characterized NADs in mouse embryonic stem cells (mESC), a tractable system for studying how NADs change during differentiation. As in MEFs (Vertii et al. 2019), we identified a large subset of mESC NADs that overlap with LADs (Type I NADs), and a smaller subset of NADs that do not overlap LADs (Type II NADs). However, Type II NADs are less prevalent in mESCs than in MEFs. mESC NADs are also notably less enriched in H3K27me3 modifications. Comparisons of MEF and mESC NADs also revealed enrichment of developmentally regulated genes in cell type-specific NADs. These analyses will facilitate future studies of genome dynamics during stem cell differentiation.

## Results

### Isolation of nucleoli from crosslinked F121-9 mESCs

We isolated nucleoli from formaldehyde-crosslinked hybrid F121-9 mES cells using methods previously shown to yield reproducible data using MEF cells (Vertii et al. 2019). In those studies, crosslinked and non-crosslinked MEFs were directly compared, and shown to yield highly overlapping results, with crosslinked samples detecting a greater proportion of the genome associated with nucleoli (Vertii et al. 2019). This suggests crosslinking could assist detection of weak or transient nucleolar interactions. Therefore, we used crosslinking for all nucleoli isolation experiments here (Fig. 1A). The purity of isolated nucleoli was confirmed using phase-contrast microscopy (Fig. 1B). Immunoblot analysis of nucleolar fractions showed that they were enriched for nucleolar protein fibrillarin relative to beta-actin (Fig. 1C). Quantitative PCR analysis revealed 9-18-fold enrichment of 45S rDNA sequences in purified nucleolar DNA relative to genomic DNA (Fig 1D). These results indicated the enrichment of nucleoli in our preparations, hence we proceeded with whole-genome sequencing of nucleolar DNA.

**Figure 1.**
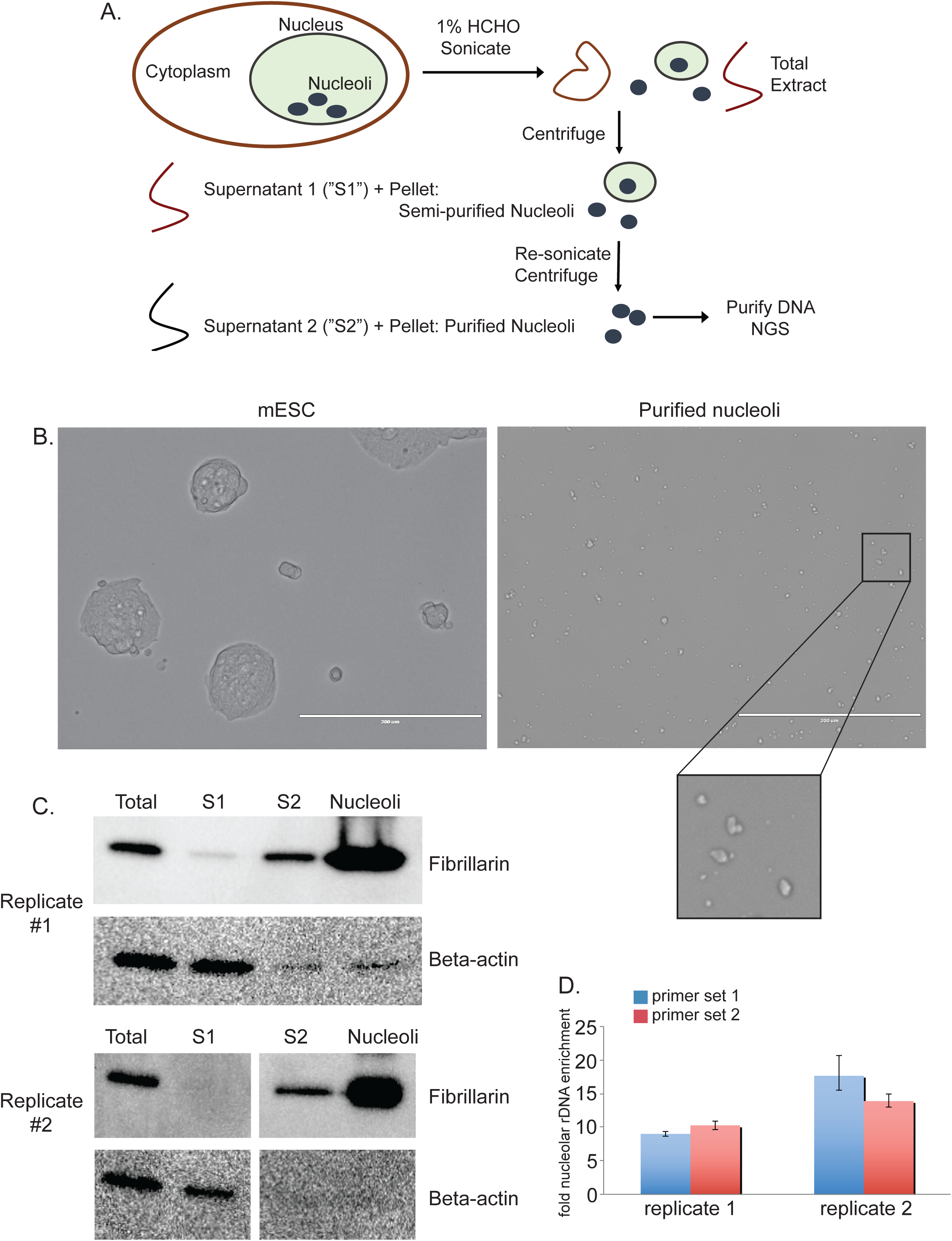
Isolation and characterization of purified nucleoli in mESC. A. Schematic diagram of nucleoli isolation from crosslinked cells. B. Phase-contrast microscopy images of F121-9 mESC grown in colonies (left panel), and nucleoli purified from them (right panel). 20x magnification, scale bar 200 μm. The inset (lower right) shows a 3x magnified image of the purified nucleoli. C. Immunoblots of fractions generated during nucleoli isolation from two replicate experiments. Fractions are labeled as shown in Fig. 1A. Fibrillarin was enriched, and beta-actin depleted, in nucleolar fractions. D. RT-qPCR measurement of 45S rDNA enrichment in nucleolar DNA from replicate experiments 1 and 2. Two different primer sets were used. Data are represented as mean enrichment relative to genomic DNA, error bars represent standard deviations for triplicate technical measurements.

### Bioinformatic analysis of NADs

We performed two biological replicate preparations of crosslinked F121-9 mESC nucleoli. In each replicate experiment, we extracted nucleolar-associated DNA from nucleoli, along with genomic DNA from whole cells from the same population of cells. We sequenced approximately 50 million reads from each nucleolar and genomic DNA sample. We note that subsampling analyses of larger MEF datasets previously showed that the number of peaks detected had reached a plateau at this sequencing depth (Vertii et al. 2019). Genomic reads were mostly uniformly distributed across the genome, whereas nucleolar reads contained well-defined peaks and valleys, with peaks overlapping known heterochromatic regions, such as constitutive LADs (cLADs) (Peric-Hupkes et al. 2010) and late replicating regions (Hiratani et al. 2010) (Fig 2A, B). cLADs were previously defined as LADs that are lamina-associated in mESCs, and also in neural precursor cells (NPCs) and astrocytes differentiated from these mESCs (Peric-Hupkes et al. 2010). Previous studies of NADs have identified frequent overlap of NADs with LADs (van Koningsbruggen et al. 2010; Németh et al. 2010; Dillinger et al. 2017; Vertii et al. 2019) and with late-replicating regions (Dillinger et al. 2017; Vertii et al. 2019), thus we concluded that the nucleolar reads are enriched with bona fide nucleolar heterochromatic regions in F121-9 mESCs.

**Figure 2.**
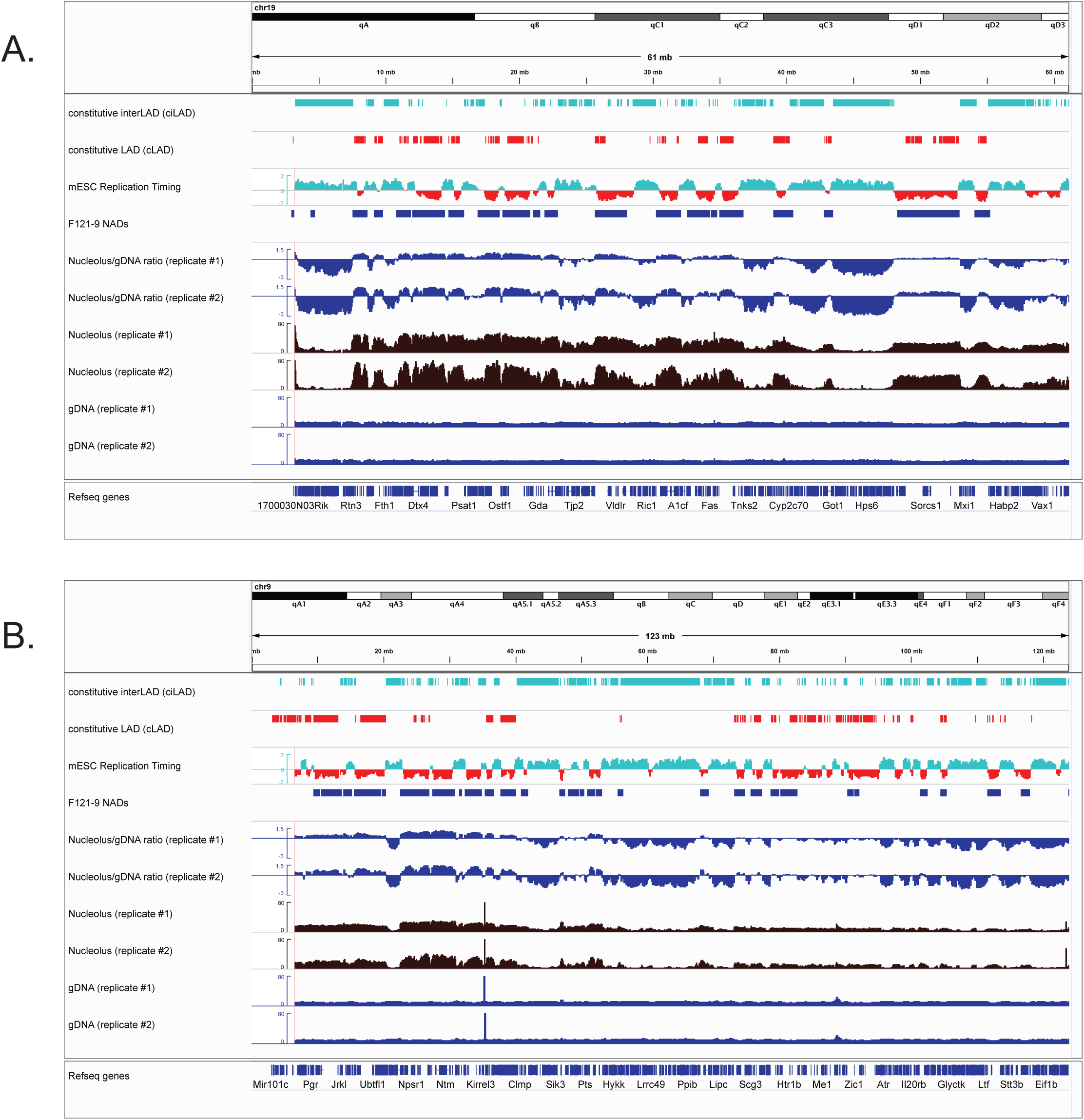
Analysis of F121-9 NAD sequencing data and comparison with heterochromatin. A. All of chromosome 19 is shown, which contains strongly nucleoli-associated regions. From the top, tracks shown are: Constitutive interLADs (ciLADs, cyan) and Constitutive LADs (cLADs, red) (Peric-Hupkes et al. 2010); mESC replication timing (Hiratani et al. 2010, early replicating regions in cyan and late replicating regions in red); F121-9 cell NAD peaks (“F121-9 NADs”, called using *NADfinder* software based on two replicate experiments); Nucleolar/gDNA ratios, shown for both replicate experiments; raw read counts from both replicates for nucleoli-associated (“Nucleolus”, brown) DNA and total genomic DNA (“gDNA”, dark blue). B. As in panel A, with all of chromosome 9 shown.

Calculating the log ratio of nucleolar reads to genomic reads resulted in a raw metric of nucleolar association across the genome (Nucleolus/gDNA ratio tracks in Fig. 2A, B). As in MEFs, visual inspection of the nucleolus/genomic ratio in mESC revealed a negative slope across chromosomes, especially noticeable on large chromosomes (Fig. 2B). Mouse chromosomes are acrocentric, i.e. the centromere is found at one end of a chromosome, and by convention these are annotated on the left. Because pericentromeric regions frequently associate with nucleolar periphery (Ragoczy et al. 2014), nucleolar associations on centromeric end of chromosomes are usually more frequent. As we have demonstrated previously using MEFs data, peak calling based only on nucleolar/genomic ratio would result in identifying peaks mostly at the centromeric end and missing the smaller peaks at the end of chromosome distal to the centromere. For this reason, we used our previously described Bioconductor package named *NADfinder* (Vertii et al. 2019) to call NAD peaks in F121-9 mESCs. This software uses local background correction, which was important for detection of validated NAD peaks distal from centromeres in MEFs (Vertii et al. 2019). *NADfinder* peak calling was performed using the default settings with a 50kb window size, a testing threshold of log2(1.5) for background corrected log2(nucleolar/genomic) ratio to define the null hypothesis, and adjusted p-value < 0.05 (Vertii et al. 2019). Potential peaks were further filtered to be > 50 kb long and to have log2 ratio > 1.7.

### 3D immuno-FISH confirmation of NAD peaks in F121-9 mESCs

To validate associations of NADs with nucleoli by an orthogonal method, we performed 3D immuno-FISH experiments, scoring association of BAC DNA probes with nucleolar marker protein fibrillarin (Figs. 3-4). We tested the association of a euchromatic negative control probe, pPK871, which lacks nucleolar association in MEFs (Vertii et al., 2019) and did not contain a peak in our F121-9 NAD-seq data. The frequency of nucleolar association for this probe was ∼24% (Fig. 4A, B). Three additional non-NAD BAC probes (pPK825, pPK1000, and pPK1003) displayed similar levels of nucleolar association (Fig. 4A). The average association frequency for these non-NAD probes in F121-9 cells is 22%, similar to the 20% frequency observed in MEF cells (Vertii et al., 2019). These observations result from stochastic positioning of loci within the nuclear volume. We note that pPK825 was also not associated with nucleoli in MEFs, whereas pPK1000 and pPK1003 had not been tested in MEFs (Vertii et al. 2019).

**Figure 3.**
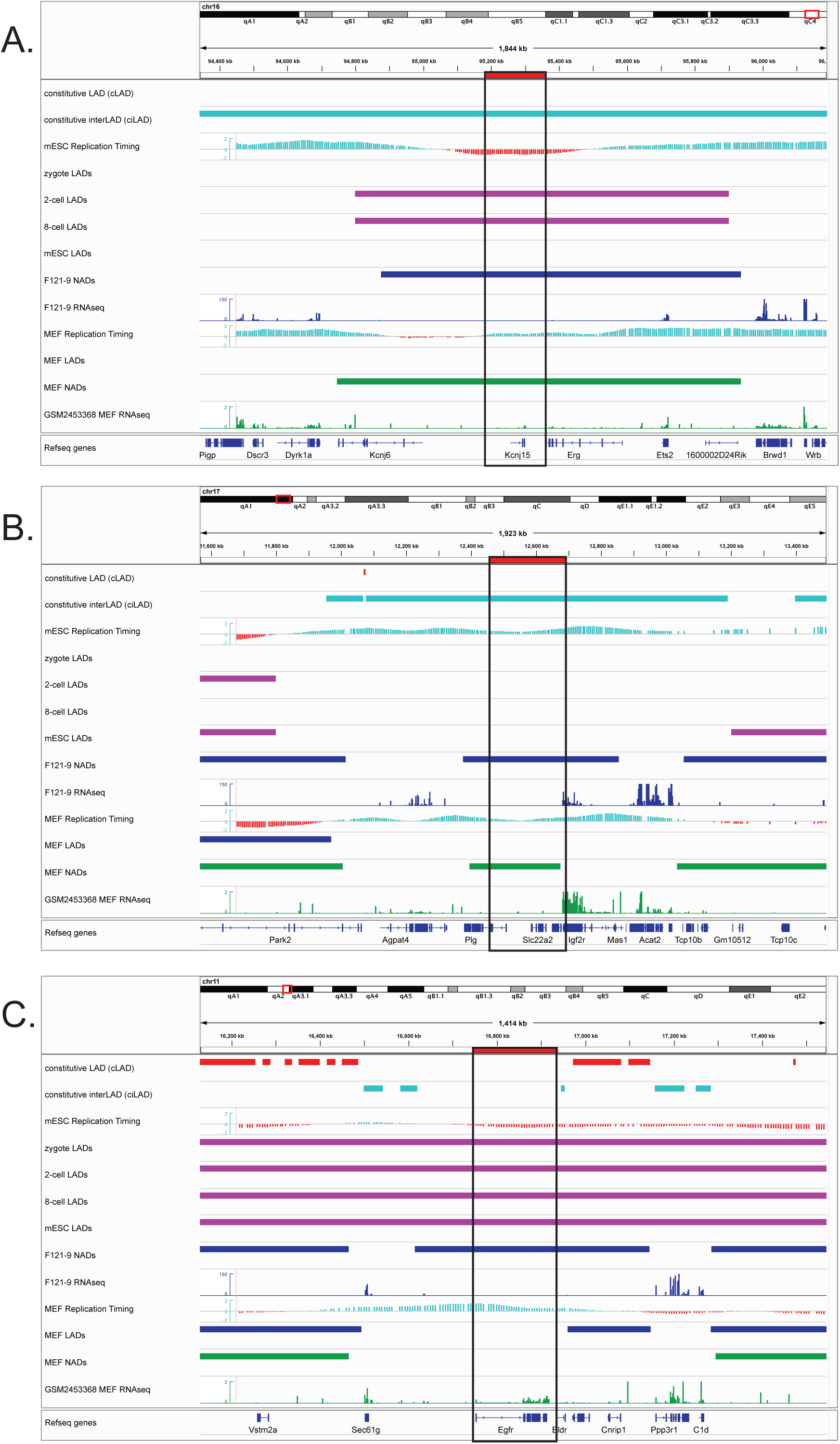
Genomic locations of BACs used for FISH experiments. For each panel, BAC locations are outlined by a black box and indicated with a red horizontal bar above the top track. From the top, tracks include cLADs (red) and ciLADs (cyan) (Peric-Hupkes et al. 2010), followed by mESC replication timing (Hiratani et al. 2010). Next are LADs from the indicated early embryonic stages (magenta) (Borsos et al. 2019), followed by F121-9 cell NAD peaks (blue) and RNA-seq data from the same preparations of F121-9 cells used to generate the NAD data. At the bottom are data from MEF cells for comparison: replication timing (Hiratani et al. 2010), LADs (Peric-Hupkes et al. 2010), NAD peaks from crosslinked cells (Vertii et al. 2019) and RNA-seq (GSM2453368 (ENCODE Project Consortium 2012)). A. pPK914. This BAC is within a NAD in both F121-9 and MEF cells, and its overlap with a ciLAD region (cyan) indicates a lack of lamina association in these cell types. However, it does become lamina-associated in the 2-cell and 8-cell stages of early embryonic development (Borsos et al. 2019). This NAD contains ion channel genes (Kcnj6, Kcnj15) and Ets-family transcription factors (Erg, Ets2). B. pPK915. This ciLAD-overlapped BAC is a NAD in both F121-9 and MEF cells, encoding solute carrier membrane transport proteins (Slc22a1, 2, 3) and plasminogen (Plg). C. pPK999. This BAC overlaps a late-replicating LAD that contains the genes encoding epidermal growth factor receptor (Egfr), EGFR Long Non-coding Downstream RNA (Eldr), pleckstrin (Plek), and cannabinoid receptor interacting protein 1 (Cnrip1). This NAD is part of a LAD throughout early embryonic development, at zygote, 2-cell, 8-cell and mESC stages (Borsos et al. 2019). Note that in MEF cells this region is not identified as a NAD, is early replicating, and displays greater expression of Egfr.

**Figure 4.**
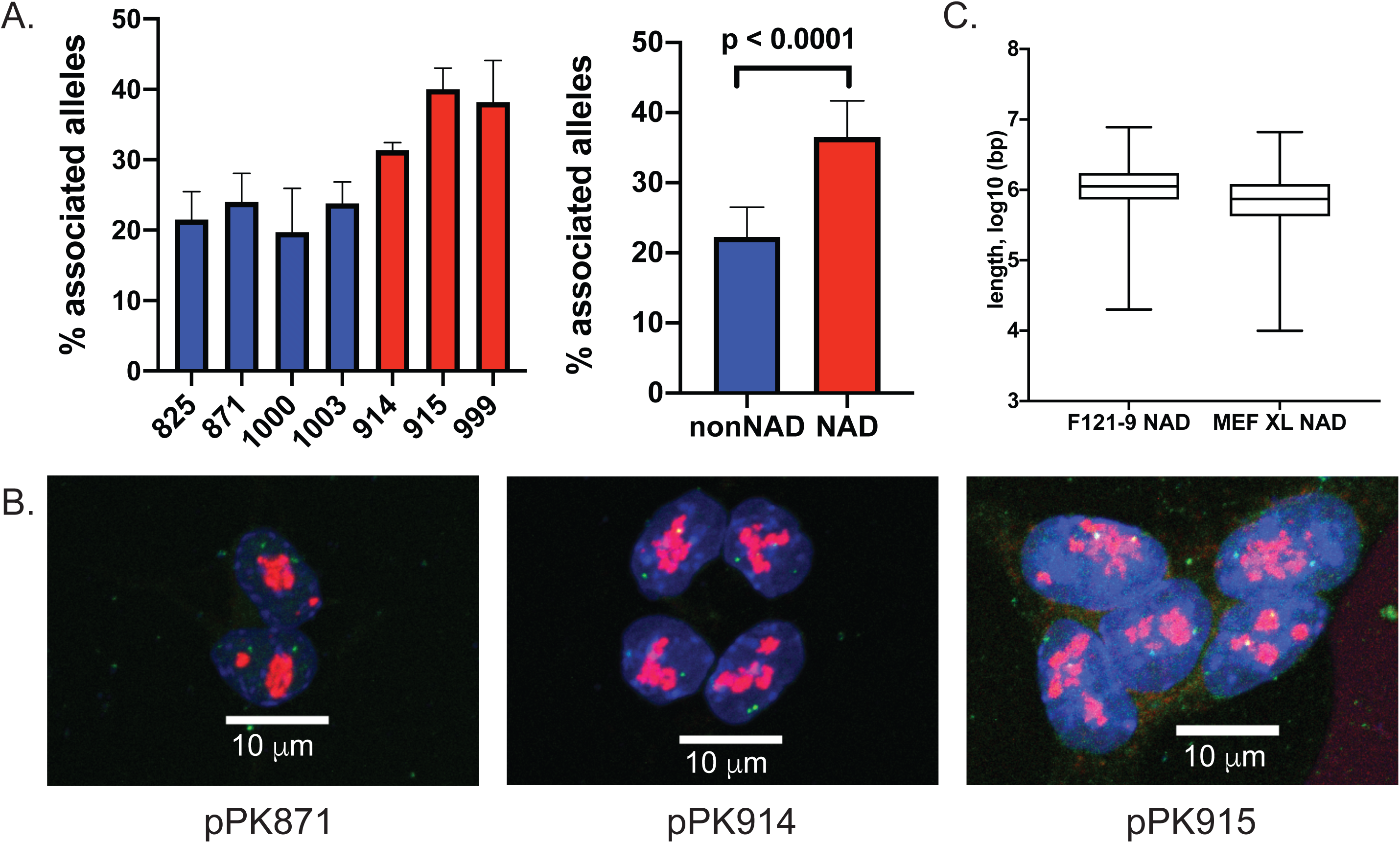
3D DNA-FISH experiments validate nucleolar association of NADs in F121-9 mESC. A. *Left*: graph of percentage of alleles that are nucleolar-associated (mean ± standard deviation for n = 3 biological replicates) for the indicated (see Supplmental Table 2) non-NAD BAC probes (blue bars) and NAD probes (red bars). *Right*: data from the left graph were grouped into non-NADs (blue bar) and NADs (red bar). NADs display significantly greater nucleolar association than non-NADs (p < 0.0001, Welch’s t-test). B. Maximum projection images from 3D immuno-FISH experiments with nuclear DAPI staining in blue, anti-fibrillarin antibody staining in red, and DNA probes (pPK871, pPK914 and pPK915) in green. 63x magnification, scale bar 10 μm. C. Length distribution of F121-9 NADs, compared to those from crosslinked MEF cells (Vertii et al. 2019).

We also analyzed BAC probes pPK914 and pPK915 (Fig. 3A, B), which overlap NAD peaks in both our F121-9 data and in MEFs (Vertii et al. 2019). In F121-9 cells, we observed that both of these probes displayed more frequent nucleolar association than did the the set of non-NAD probes (pPK914, p < 0.0001; pPK915, p = 0.0002, Welch’s t-test), indicating that these regions are NADs in both MEFs and F121-9 mESCs (Fig. 4A, B). Both the pPK914 and pPK915 probes overlap ciLAD regions, which means that these regions were not observed to associate with lamina in mESCs or MEFs (Peric-Hupkes et al. 2010). However, recent LAD maps of early mouse embryogenesis (Borsos et al. 2019) show that pPK914 probe is lamina-associated in 2-cell and 8-cell embryos (Fig. 3A). Therefore, this region is nucleolar-associated in both mESCs and somatic cells, but lamina-associated only during limited periods in very early development. We also analyzed a region detected as a NAD in mESCs, but not in MEFs (pPK999, Fig. 3C). FISH analysis showed that this probe indeed displayed increased nucleolar association compared to non-NAD probes in F121-9 cells (Fig. 4A; p = 0.0220, Welch’s t-test). We note that this probe is lamina-associated throughout early embryonic stages (zygote, 2-cell, 8-cell embryos and mESCs), but not in somatic MEF cells (Fig. 3C). Furthermore, pPK999 contains the *Egfr* gene, for which transcript levels are higher in MEFs (FPKM value 51.5) (Delbarre et al. 2017) compared to mESCs (FPKM value 0.2 (Supplemental Table 1)). This is an example of a genomic locus that is nucleolar-associated and transcriptionally repressed in mESCs, and which is no longer associated and becomes more active in MEFs. In sum, these FISH data demonstrate that the identified NADs include bona fide nucleolar heterochromatic regions in F121-9 mESCs, conserved or regulated during cell differentiation.

The length of F121-9 NADs ranges up to 8 Mb (Fig. 4C), with median length 1.1 Mb, which is slightly larger than median length of MEF NADs, 0.7 Mb (Vertii et al. 2019). We noted that NADs in F121-9 cells covered 31% of the non-repetitive genome, a smaller percentage than observed in crosslinked MEF NADs (41%) (Vertii et al. 2019). The 31% fraction of the mESC genome in NADs is also smaller than the fraction of the mouse genome in LADs, either for embryonic stem cells or somatic cells (∼40%) (Peric-Hupkes et al. 2010), or during early mouse embryogenesis (∼40-60%) (Borsos et al. 2019) (see Discussion).

### Two types of NADs in F121-9 mESCs

In our previous analysis of MEF data, we had defined a “Type I” class of NADs as those overlapping LADs (Vertii et al. 2019). Additionally, a contrasting “Type II” class of NADs was defined which overlaps “constitutive interLADs” (ciLADs), the regions defined as those which were not lamina-associated during multiple steps of cellular differentiation (Peric-Hupkes et al. 2010). In MEFs, Type I NADs are approximately five-fold more abundant, and tend to replicate late; in contrast, the less abundant Type II NADs more frequently overlap with early replicating regions (Vertii et al. 2019). In F121-9 mESC NADs, we also observed abundant Type I NADs that overlap with cLADs (421 Mb of the total 845Mb NAD population; Fig. 5A). However, Type II NADs that overlap with ciLADs comprise only 77 Mb, much less than the 147 Mb observed in similarly crosslinked MEFs (Fig. 5A; Vertii et al. 2019). Visual inspection of the distribution of the two classes in a genome browser illustrated the greater size of the Type I subset compared to Type II regions (Fig. 5B). Despite the small size of the F121-9 Type II NAD subset, we note that we have validated nucleolar association of two Type II NAD probes (pPK914, pPK915; Fig. 4A, B). These two probes have previously been confirmed to lack significant lamina association in MEFs (Vertii et al. 2019). Both overlap ciLAD regions (Fig. 3A, B), indicating that they lack lamina association during multiple steps in the process of differentiation from mES cells to astrocytes (Peric-Hupkes et al. 2010; Meuleman et al. 2013). We conclude that in mES cells, as in MEFs, a large proportion of NADs overlap LAD regions, but that the amount of ciLAD overlap in mES cells is smaller.

**Figure 5.**
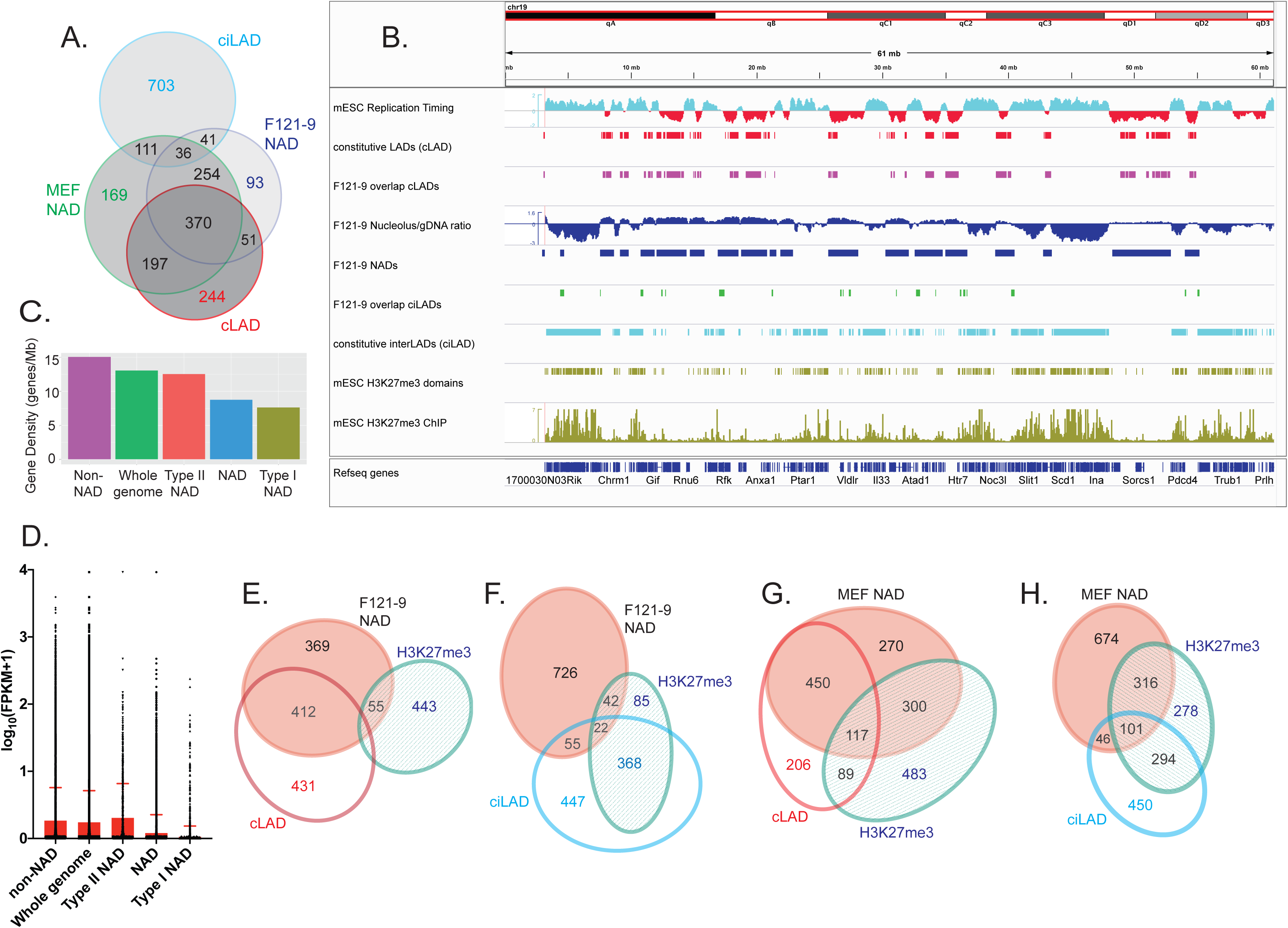
Two types of NADs in F121-9 mESC. A. Venn diagram illustrating the overlaps among F121-9 NADs, MEF NADs (Vertii et al. 2019), cLAD, and ciLAD regions (Peric-Hupkes et al. 2010). Numbers show the size of the indicated regions in Mb. B. Chromosomal view of F121-9 NADs overlapping cLADs and ciLADs. The entire chromosome 19 is shown. Euchromatic features (early replication timing, ciLAD) are displayed in cyan, and heterochromatic features (late replication timing, cLAD) are shown in red. From the top, displayed tracks are mESC replication timing (Hiratani et al. 2010), cLAD (Peric-Hupkes et al. 2010), NAD overlap with cLAD (i.e. Type I NADs, magenta), nucleolar/genomic ratio and NAD peaks (blue), NAD overlap with ciLAD (i.e. Type II NADs, green), ciLAD (Peric-Hupkes et al. 2010), H3K27me3 domains, and mESC H3K27me3 ChIP-seq data (Cruz-Molina et al. 2017) used for H3K27me3 domain identification (olive green). C. Gene densities (genes/Mb) of the indicated regions, ranked left to right. “NAD” indicates all F121-9 NADs. D. A box plot of gene expression levels from F121-9 RNA-seq data, expressed as log_10_(FPKM+1) for the same indicated genomic regions as in panel C. The top of the red box indicates the mean value for each population, and the standard deviation is marked by the red error bar. E. Venn diagram illustrating the overlaps among F121-9 NADs, cLADs (Peric-Hupkes et al. 2010) and mESC H3K27me3 domains (Cruz-Molina et al. 2017). Numbers indicate the size of regions in Mb. The overlaps among all three sets (9 Mb) and between the cLAD and H3K27me3 sets (10 Mb) are left off the diagram because of their small sizes. Diagram was generated using eulerAPE 3.0. F. As in panel E, except here the overlap analysis includes ciLADs (Peric-Hupkes et al. 2010) instead of cLADs. G. As in panel E, except here Venn diagram illustrates the overlaps among crosslinked MEF NADs (Vertii et al. 2019), cLADs (Peric-Hupkes et al. 2010) and MEF H3K27me3 domains (Delbarre et al. 2017). H. As in panel G, except here the overlap analysis includes ciLADs (Peric-Hupkes et al. 2010) instead of cLADs.

We then analyzed gene density and gene expression characteristics of the different NAD subsets from F121-9 cells. As we had observed in MEFs (Vertii et al., 2019), gene density of Type II NADs was greater than that of NADs as a whole, which in turn have higher gene density compared to Type I NADs (Fig. 5C). Using RNA-seq data we obtained from the same preparations of F121-9 cells that were used for nucleolar purification, we analyzed genomic trends in steady-state mRNA levels by plotting the distributions of the FPKM values. As in MEFs (Vertii et al. 2019), F121-9 NADs displayed lower FPKM values than the genome-wide average (p < 0.0001). In addition, FPKM values for the Type I NAD subset were significantly lower than those for NADs as a whole (p < 0.0001) (Fig. 5D). Thus, Type I NADs in both MEFs and F121-9 cells display low gene expression levels characteristic of heterochromatin. In contrast, in F121-9 cells Type II NADs displayed mean gene expression levels that are slightly higher than those observed in the whole genome (p < 0.0003) or even in non-NAD regions (p < 0.0233) (Fig. 5D). Therefore, in both F121-9 cells and MEFs (Vertii et al., 2019), Type II NADs can become associated with nucleoli without adopting the highly silenced status of Type I NADs.

However, F121-9 NADs displayed a prominent difference from those in MEFs, regarding overlap with H3K27me3 peaks. We note that H3K27me3 is functionally important for heterochromatin localization because Ezh2 inhibitors that block this modification decrease laminar and nucleolar associations by heterochromatin (Harr et al. 2015; Vertii et al. 2019). In MEFs, we observe frequent overlap of H3K27me3 peaks (Delbarre et al. 2017) with both Type I (117 Mb out of 567 Mb) and Type II NADs (101 Mb out of 147 Mb) (Fig. 5G, H; Vertii et al. 2019). In contrast, in F121-9 cells we observed that overlap of NADs with H3K27me3-enriched domains (Cruz-Molina et al. 2017) was much smaller than observed in MEFs: only 9 Mb of the 421 Mb of Type I NADs and 22 Mb of 77 Mb of Type II NADs overlap with H3K27me3 domains (Fig. 5E, F). These differences likely reflect the lower abundance of repressive histone marks in mESCs compared to differentiated cells; this includes H3K27me3, which becomes more abundant during differentiation ((Martens et al. 2005; Hawkins et al. 2010; Atlasi and Stunnenberg 2017); see Discussion). Indeed, our analysis of an F121-9 data set (Cruz-Molina et al. 2017; see Methods) detected 517 Mb of H3K27me3 peak regions in F121 cells, and an almost two-fold larger amount (990 Mb) was found in MEFs (GSM1621022; Delbarre et al. 2017)). However, we note that the amount of H3K27me3 peaks in NADs is much more than two-fold greater in MEFs (417 Mb, Fig. 5G, H) than in F121-9 cells (66 Mb, Fig. 5E,F). Together, these data suggest that H3K27 methylation is a key aspect of NAD chromatin maturation that has not yet occurred fully in mES cells (see Discussion).

### Cell type-specific and conserved NADs

We compared F121-9 stem cell NADs with crosslinked MEF NADs (Vertii et al. 2019), defining overlapped regions on a nucleotide-by-nucleotide basis (e.g. Fig. 6A). Close to 80% (660 Mb) of nucleotides in stem cell NADs overlap with nucleotides in MEF NADs (Fig. 5A). We designate NADs shared by MEFs and F121-9 stem cells as “conserved NADs”. Analysis of the intersection of conserved NADs with cLAD and ciLAD regions revealed that more than half of conserved NADs overlap cLADs (370 Mb; Fig. 5A), which are the most gene-poor subset of LADs and are generally poorly expressed, constitutive heterochromatin (Peric-Hupkes et al. 2010; Meuleman et al. 2013; van Steensel and Belmont 2017). Consistent with these trends, Jaccard similarity coefficient analysis indicated high correlation of conserved NADs with cLADs and late replicating regions (Marchal et al. 2018) (Fig. 6B). Furthermore, the conserved NADs display the lowest transcript levels in both cell types (Fig. 6C-F), as expected due to the constitutive heterochromatic features of these regions.

**Figure 6.**
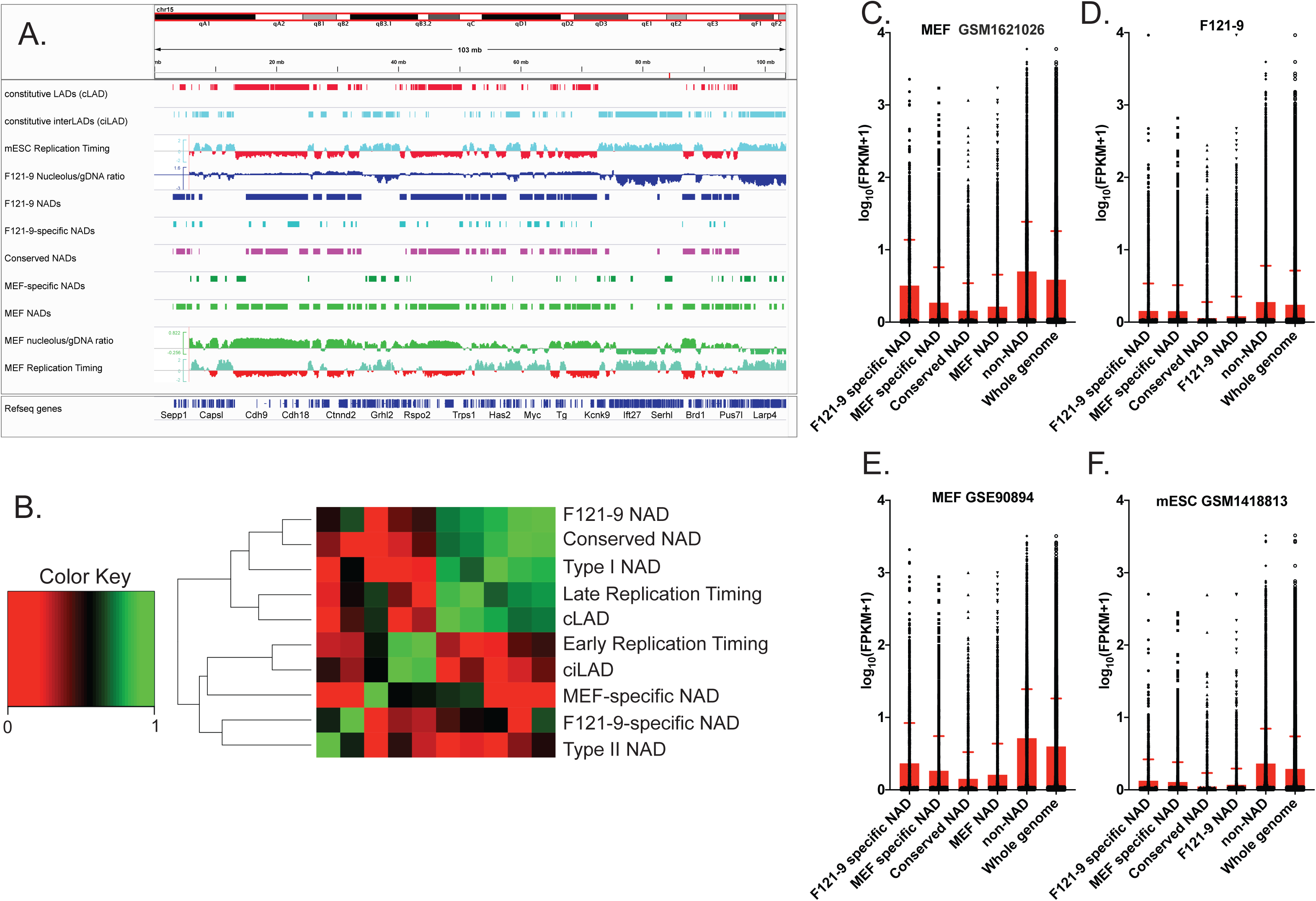
Conserved and cell type-specific NADs. A. IGV browser view of entire chromosome 15. Euchromatic features (early replication timing, ciLAD) are displayed in cyan, and heterochromatic features (late replication timing, cLAD) are shown in red. From the top, tracks shown are cLAD, ciLAD (Peric-Hupkes et al. 2010), mESC replication timing (Hiratani et al. 2010), F121-9 nucleolar/genomic ratio and F121-9 NAD peaks (blue), “F121-9 specific NADs”, i.e. NADs found only in F121-9 cells (light blue), “conserved NADs”, or NADs shared between F121-9 and MEFs (magenta), “MEF-specific NADs” (dark green), MEF NAD peaks and MEF nucleolar/genomic ratio (Vertii et al. 2019) in green, and MEF replication timing (Hiratani et al. 2010). B. Jaccard similarity coefficients were grouped based on similarities among the indicated regions. “F121-9 NAD” indicates all NADs identified in F121-9 cells in this study. “Conserved NAD” indicates NADs shared between F121-9 and MEF NADs (Vertii et al. 2019), whereas “F121-9-specific NAD” indicates NADs detected in F121-9, but not MEF cells. Conversely, “MEF-specific NAD” indicates NADs found in MEFs, but not in F121-9 cells. “Type I NAD” indicates F121-9 NADs that overlap with cLADs, and “Type II NAD” indicates F121-9 NADs that overlap with ciLADs (Peric-Hupkes et al. 2010). “cLAD” and “ciLAD” regions are from Peric-Hupkes et al. 2010, and F121-9 early replication timing and late replication timing regions are from Marchal et al. 2018. Note that F121-9 NADs, conserved F121-9 NADs, cLADs and Type I NADs are highly similar. In contrast, Type II NADs are most similar to F121-9-specific NADs. C. A box plot of gene expression levels from MEF RNA-seq data (GSM1621026; Delbarre et al. 2017), expressed as log_10_(FPKM+1) for the indicated subsets of NAD, non-NAD and whole genome regions. The statistical significance of pairwise comparisons were all p < 0.0001 (Welch’s t-test). D. As in panel C, except our F121-9 RNA-seq data is used for FPKM analysis. The indicated pairwise comparisons were all statistically significant (p < 0.0001), except for that between F121-9 and MEF-specific NADs do not achieve statistical significance (p = 0.82). E. As in panel C, except different MEF RNA-seq data (GSE90894; (Chronis et al. 2017)) was used for FPKM analysis. The changes between cell type-specific NADs achieve statistical significance (p<0.0001, Welch’s t-test). F. As in panel C, except mESC RNA-seq data (GSM1418813; (Lowe et al. 2015)) is used for FPKM analysis. The changes between F121-9 and MEF-specific NADs do not achieve statistical significance (p=0.13).

We next turned our attention to NADs found only in one of the two analyzed cell types. The Jaccard analysis indicated that these cell type-specific NAD regions (i.e. “MEF-specific NADs” and “F121-9-specific NADs”) are distinct from the conserved NADs, clustering separately from conserved NADs, cLAD and late replicating regions (Fig. 6B). We analyzed steady-state mRNA levels in conserved and cell type-specific NADs by using FPKM values from F121-9 and MEF (Delbarre et al. 2017) RNA-seq data (Fig. 6C, D). As we expected, MEF RNA-seq data revealed lower levels of transcripts from genes within MEF-specific NADs than from F121-9-specific NADs (p-value < 0.0001) (Fig. 6C), indicating that in MEFs, nucleolar association correlates with transcriptional silencing. In contrast, our RNAseq data from F121-9 cells showed that transcript levels within both the MEF-specific NADs and the F121-9-specific NADs are statistically indistinguishable (p-value = 0.82) (Fig. 6D). We observed similar trends in independent sets of MEF and mESC RNA-seq data from the literature (Lowe et al. 2015; Chronis et al. 2017) (Fig. 6 E, F). These observations were unexpected in that the MEF-specific NADs are not nucleolar-associated in the F121-9 cells, yet are on average less highly expressed than non-NAD genes in these cells. These data suggest that in F121-9 stem cells, gene repression could precede localization to the nucleolar periphery that occurs later during cellular differentiation (see Discussion).

### Gene Ontology analysis of conserved and cell-type specific NADs

To further characterize the conserved NADs, we next analyzed enriched GO-terms within these. The most significantly enriched Molecular Functions term was “Response to smell detection” (Fig. 7A; Supplemental Table 3), including olfactory receptor (OR) and vomeronasal receptor genes. These clustered genes are not expressed in either stem cells or fibroblasts and are frequently within NADs in both F121-9 stem cells and MEFs (e.g. the OR genes on chr11, Fig. 7B). Among other well-represented gene families in conserved NADs were cytochrome P450 family members: *Cyp2a12, Cyp2b10, Cyp2c50* (“heme-interacting genes” in Fig. 7A), which are responsible for breaking down toxins, as well as synthesizing steroid hormones, fats and acids, and are most highly expressed in liver (Hannemann et al. 2007). Neurotransmitter receptors were also enriched for conserved NADs, for example, genes that encode for glutamate receptors (*Gria2, Grid2*, etc.), GABA-A receptors (*Gabra5, Gabrb1*, etc.) and glycine receptors (*Glra1, Glrb*, etc.). The common thread among these gene classes is in that they are developmentally regulated, and most strongly induced in lineages not represented by embryonic stem cells or fibroblasts.

**Fig. 7.**
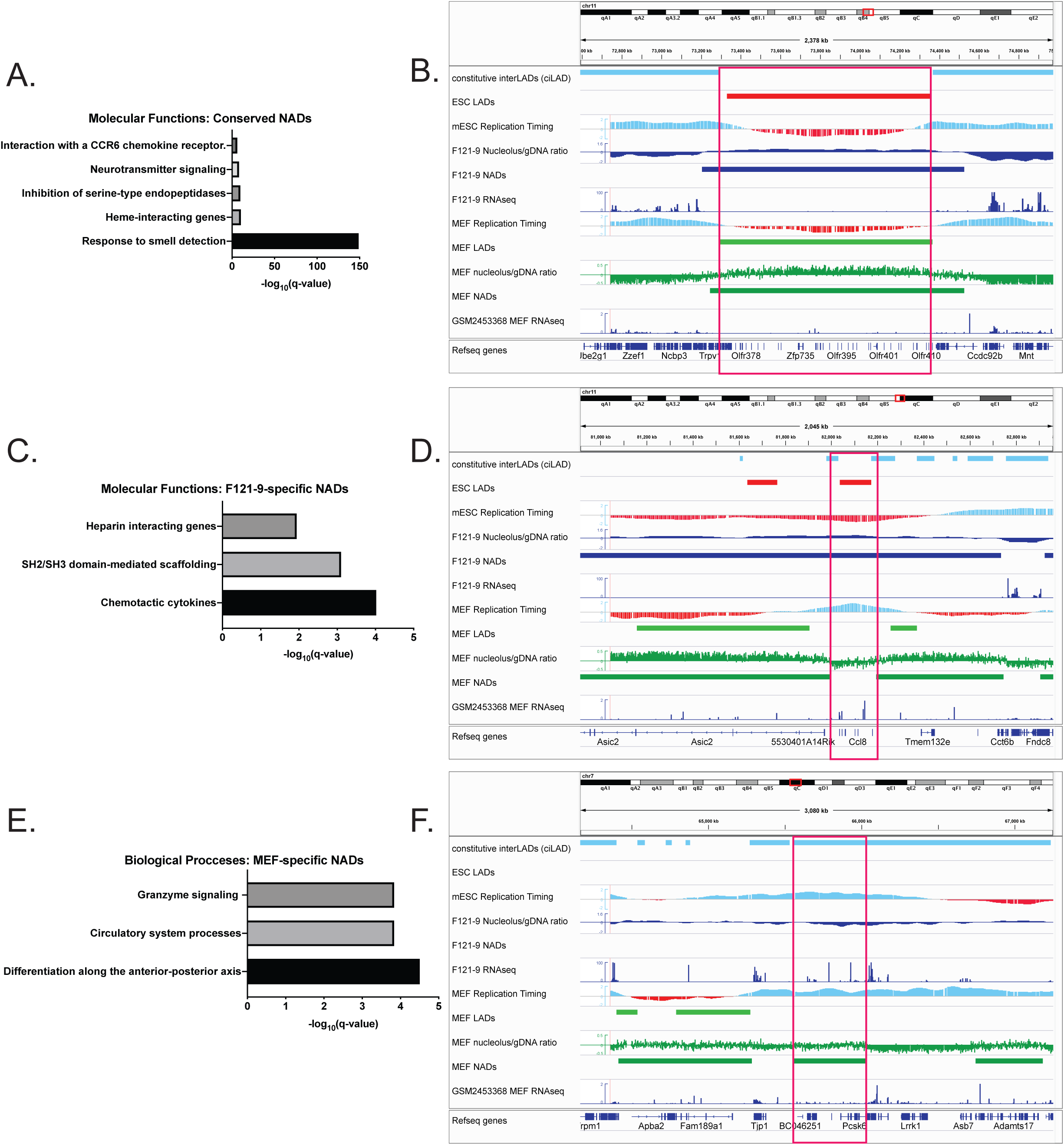
GO analysis of conserved and cell type-specific NADs. A. Molecular Functions subset of GO enrichment analysis of conserved NADs, with −log_10_(q-values) shown. B. Genomic region containing NAD peak (red box) conserved in both MEF and F121-9 cells. This peak contains a cluster of olfactory genes on chromosome 11. ciLAD, mESC and MEF replication timing tracks are displayed as in Fig. 5B. The other tracks shown from the top are mESC LADs (Peric-Hupkes et al.2010; red), F121-9 nucleolar/genomic ratio, NADs and RNA-seq data (blue), MEF LADs (Peric-Hupkes et al. 2010), MEF nucleolar/genomic ratio, NADs (Vertii et al. 2019) (green) and RNA-seq (GSM2453368 (ENCODE Project Consortium 2012)) (blue). C. Molecular Functions subset of GO enrichment analysis of F121-9-specific NADs. D. As in panel B, showing genomic region corresponding to F121-9-specific NAD (red box), overlapping *Ccl* family of chemokine ligands. E. Biological Functions subset of GO enrichment analysis of MEF-specific NADs. F. As in panel B, showing genomic region containing MEF-specific NAD (red box), overlapping the *Pcsk6* gene important for differentiation along anterior-posterior axis.

We next analyzed the F121-9-specific NADs. Among these, chemotactic cytokines were the GO-derived “Molecular Functions” class with the lowest q-value (Fig. 7C; Supplemental Table 4). The majority of these chemokines are represented by the CC chemokine ligand family, a cluster of which is shown in Fig. 7D. This cluster of *Ccl2, Ccl12 and Ccl1* genes has heterochromatic features in the F121-9 cells: late replication timing, no steady-state mRNA transcripts, presence within both LAD and NAD regions. In contrast, in MEFs this gene cluster is within neither NAD nor LAD sequences and has euchromatic features, including early replication timing and high gene transcript levels. This is an example of a genomic region in which multiple features are altered, becoming more euchromatic upon differentiation.

We then considered the converse case, the MEF-specific NADs. Among these, the “Biological Processes” GO classifications included genes responsible for differentiation along the anterior-posterior axis (Fig. 7E; Supplemental Table 5), an example of which is *Pcska6* gene (Fig. 7F). This genomic region displays euchromatic features (overlapping a ciLAD region, early replicating timing and high transcript levels: FPKM value 22.2) in mESCs, befitting the need for anterior-posterior axis establishment factors at this early developmental stage. In MEFs, this locus displays altered features, becoming nucleolar-associated, and generating reduced transcript levels (FPKM value 6.6) (Delbarre et al. 2017). In general, both conserved and cell type-specific NADs generally include genes that display reduced expression levels, suggesting that nucleolar localization could contribute to (or be a consequence of) the transcriptional silencing of resident genes. A major question remains as to how functionally distinct classes of NADs (e.g. Type I and Type II NADs) are targeted to nucleoli, and how this has distinct transcriptional consequences in each case (e.g. Fig. 5D; see Discussion).

## Discussion

### Heterochromatin formation during differentiation

Several types of evidence indicate that compared to differentiated cells, chromatin in mESCs is less condensed, and the ratio of euchromatin to heterochromatin is higher (Gaspar-Maia et al. 2011). For example, fluorescence recovery after photobleaching experiments demonstrated that mESCs display more highly mobile core and linker histones, as well as Heterochromatin Protein 1 (HP1α) than do differentiated cells. These features are thought to contribute to the transcriptional hyperactivity in pluripotent stem cells (Meshorer et al. 2006; Bhattacharya et al. 2009). For example, many repetitive elements that are silent in somatic cells are transcribed in mESCs (Efroni et al. 2008). Microscopy studies showed that electron-dense heterochromatic structures are less condensed and less frequently localize near nuclear lamina in mESCs compared to heterochromatin in differentiated cells (Hiratani et al. 2010; Ahmed et al. 2010; Mattout et al. 2015). Particularly relevant to our studies, more prominent electron-dense perinucleolar heterochromatin-like structures have been observed in differentiated cells, such as NPCs, compared to mESCs (Savić et al. 2014). In concert with changes in the appearance and localization of heterochromatin, the abundance of heterochromatic marks such as H3K27me3, and H3K9me3 increases during differentiation (Lee et al. 2004; Martens et al. 2005; Meshorer et al. 2006; Wen et al. 2009; Hawkins et al. 2010). Together, these data are consistent with our observation that NADs in mESCs comprise a smaller fraction of the genome compared to MEFs (31 vs. 41%). Likewise, genome coverage by LADs increases during differentiation. For example, a recent study shows that LADs are first established immediately after fertilization, preceding TAD formation and instructing A/B compartment establishment (Borsos et al. 2019).

### The Type II class of NADs is different in stem cells and fibroblasts

Two functionally distinct classes of NADs have recently been reported in mouse embryonic fibroblasts (Vertii et al. 2019). Here, we show that in F121-9 mESCs, Type I NADs that overlap LAD regions are frequently the same as those found in MEFs (Fig. 5A), and exhibit similar low gene expression levels as expected for constitutive heterochromatin (Fig. 5D). In contrast, the Type II NADs defined by their overlap with ciLAD regions is much smaller in F121-9 than in MEF cells (Fig. 5A). We also note that NADs in F121-9 cells display much less overlap with H3K27me3 peaks than do MEF NADs (Fig. 5E-H). Together, these data suggest that acquisition of H3K27me3, the hallmark of facultative heterochromatin (Trojer and Reinberg 2007) by NADs is part of the process of cellular differentiation. Indeed, we note that GO analysis of MEF Type II NADs showed enrichment for developmentally regulated GO terms, for example, organ morphogenesis and sensory organ development (Vertii et al. 2019). Thus, stem cells prevent developmentally important genes from acquiring characteristics of facultative heterochromatin including nucleolar association, whereas these genes can become NADs after they are no longer required during development.

### How are NADs targeted to nucleoli?

The precise mechanisms for targeting the two distinct classes of NADs to nucleoli remain unclear. Several studies implicate phase separation in the formation of heterochromatin domains (Larson et al. 2017; Strom et al. 2017; Shin et al. 2018) and nuclear bodies, such as nucleoli (Brangwynne et al.; Feric et al. 2016; Mitrea et al. 2016). Our recent data suggest that Type II NADs are more sensitive than Type I NADs to hexanediol treatment (Vertii et al. 2019). Hexanediol perturbs phase separation, likely due to interfering with weak hydrophobic interactions that are important for liquid-like condensate formation (Ribbeck and Görlich 2002). Liquid-liquid demixing reactions frequently involve proteins that have intrinsically disordered regions (IDR) and RNA recognition motifs (Feric et al. 2016), as found for example in nucleolar proteins fibrillarin (Fbl) and nucleophosmin (Npm1). Notably, depletion of Nlp, the *Drosophila* homolog of Npm1, led to declustering of centromeres and decreased association of centromeres with nucleolar periphery (Padeken et al. 2013). Therefore, it is possible that Type II NADs are specifically targeted to nucleolar periphery through the interactions between nucleolar proteins with IDRs (e.g. Npm1) with RNA species that are yet to be identified. Additionally, Polycomb repressive complex 1 (PRC1) protein chromobox 2 (CBX2) undergoes phase separation and forms liquid-like condensates in mESCs (Tatavosian et al. 2019), and Polycomb proteins are part of the MiCee complex that together with let-7 family miRNAs confers nucleolar association to specific loci (Singh et al. 2018). Therefore, Polycomb group (PcG) proteins are good candidates for nucleolar targeting of Type II NADs via phase separation. This may be especially important during differentiation, when PcG proteins gain special importance (Aloia et al. 2013; Lavarone et al. 2019). However, inhibition of PRC2 enzymatic activity decreases both nucleolar (Singh et al. 2018; Vertii et al. 2019) and laminar heterochromatin localizations (Harr et al. 2015), making it unlikely that PRC2 can target loci to a unique destination. Additionally, nucleolar localization of the *Kcnq1* locus can occur in cells lacking functional Polycomb complexes (Fedoriw et al. 2012a), indicating that multiple mechanisms likely exist. Other candidate trans-acting factors that could specifically target genomic regions to the nucleolar periphery are the proteins Ki-67 and the p150 subunit of Chromosome Assembly Factor-1 (CAF-1) (Smith et al. 2014; Matheson and Kaufman 2017), and the Kcnq1ot1 (Mohammad et al. 2008) and Firre (Yang et al. 2015) long non-coding RNAs.

### Anomalies of MEF-specific NADs in stem cells

One question of interest is whether nucleolar association leads to, or is a consequence of, transcriptional repression. Notably, previous studies have shown that tethering of loci to the nucleolar periphery via 5S rDNA sequences results in transcriptional silencing (Fedoriw et al. 2012b), so at least in that case a causal relationship has been established.

In MEF cells, genes in the MEF-specific NADs display mean expression levels lower than genes in the F121-9-specific NADs (p < 0.0001) (Fig. 6C, E). This is the expected situation, in which genes that had been in NADs earlier in development (e.g. in stem cells) become derepressed if that localization is lost. In contrast, in F121-9 cells, genes within MEF-specific NADs showed similar transcript levels as genes within F121-9-specific NADs (p = 0.82, Fig. 6D); the same was true in other mES cells analyzed (p = 0.13, Fig. 6F). Why aren’t the MEF-specific NADs more transcriptionally active in stem cells, since they haven’t yet acquired nucleolar association? This could be due to other repressive mechanisms acting on regions within MEF-specific NADs, for example, lamina association: 40% of MEFs-specific NADs overlap with cLADs (Fig. 5A). Alternatively, additional factors contributing to transcriptional repression may precede (and perhaps contribute to) nucleolar association. Development of reagents allowing control of perinucleolar associations will be key to exploring the relationship between nucleolar localization and transcriptional repression.

## Materials and Methods

### F121-9 mESC nucleoli isolation

For each preparation, cells were grown in eleven 15-cm plates and harvested one or two days after seeding them, with total cell numbers of 3-5 × 10^8^ per preparation. One hour prior to nucleoli isolation, old cell culture medium was replaced with fresh medium. Plates grown in parallel were used for genomic DNA extraction (DNeasy Blood & Tissue kit, Qiagen), and RNA extraction (TRIzol, ThermoFisher Scientific and RNeasy mini kit, Qiagen).

*Crosslinked isolation of nucleoli* was done as described previously (Vertii et al. 2019)).

### Cell culture

F121 mouse embryonic stem cell (mESC) line is a female cell line derived from a cross between male *Castaneus* and female *129* mice in Jaenisch lab (Rasmussen et al. 1999), and F121-9 was subcloned in Gribnau lab. F121-9 cells were obtained from Gilbert lab at passage 8. The cells were grown on gelatin-coated plates and cultured in 2i medium. Accutase (EMD Millipore SF006) was used to detach cells from plates and passage into new dishes. Prior to seeding cells, dishes were coated with 0.1% gelatin (EMD Millipore, SF008) for at least 25 min at room temperature, after which gelatin was aspirated. Dishes were rinsed with DPBS (Gibco, 14190144), which was aspirated, and cells were seeded in these dishes. 2i medium was obtained as described previously (Vertii et al. 2019). Cells were passaged at 3 × 10^4^/cm^2^ density. 2X HyCryo-STEM cryopreservation medium (GE Healthcare, SR30002.02) was used to freeze cells.

### Quantitative PCR

DNA was extracted from input whole cells and purified nucleoli using DNeasy Blood & Tissue kit (Qiagen). Quantitative PCR analysis was done as outlined previously (Vertii et al. 2019).

### Antibodies

The following antibodies were used: fibrillarin (Abcam, ab5821), actin (Sigma-Aldrich, A1978) and nucleophosmin (Abcam, ab10530). Secondary antibody for immunofluorescence was Alexa 594-conjugated donkey anti-rabbit (ThermoFisher, A-21207) and Alexa 594-conjugated goat anti-mouse (ThermoFisher, A-11020). For western blots, horseradish peroxidase (HRP) anti-mouse and anti-rabbit secondary antibodies (Jackson ImmunoResearch) were used.

### Immunoblotting

Proteins from total cell lysates and purified nucleoli were analyzed as noted previously (Vertii et al. 2019).

### DNA isolation, deep sequencing, and read preprocessing and mapping

Total genomic and nucleolar DNA was purified using DNeasy Blood & Tissue kit (Qiagen). Libraries were generated using NEBNext Ultra II DNA Library Prep Kit for Illumina (New England Biolabs). The DNA was fragmented to a size of 350 bp, and these fragments were size selected with sample purification beads. 150 bp paired-end sequencing was performed using Illumina reagents. 52.1 and 51.5 million reads were obtained for two replicates of genomic samples, and 49.4 and 52.8 million reads were obtained for two replicates of nucleolar samples. >95% of nucleolar samples, and >96% of genomic samples were mappable. For more information regarding sequencing, please see the files at data.4dnucleome.org under accession numbers 4DNESXE9K9DB, 4DNESUJZ5FL2. Trimming and alignment of mapped reads to the mouse genome (mm10) was done as previously described (Vertii et al. 2019).

### RNA isolation, deep sequencing, and read preprocessing and mapping

Total RNA from two replicates of F121-9 mESC were extracted using TRIzol (ThermoFisher Scientific) and purified using RNeasy mini kit (Qiagen). Libraries were constructed using NEBNext Ultra II RNA Library Prep kit for Illumina (New England Biolabs). The mRNA was fragmented, and double-stranded cDNA library synthesized, and completed through size selection and PCR enrichment. 150 bp paired-end sequencing was achieved using Illumina HiSeq 4000 platform. 22.2 and 26.7 million reads were obtained for each of the two replicates of mESC RNA. >92% of replicate 1, and >86% of replicate 2 were mappable. For more information regarding sequencing, please see the files at data.4dnucleome.org under accession number 4DNESDHILYLU. The quality of the sequencing reads was evaluated with fastqc (0.11.5) (https://www.bioinformatics.babraham.ac.uk/projects/fastqc/ The paired-end reads were aligned to the mouse genome (ensemble GRCm38) using STAR (version 2.5.3a) with ENCODE standard options as --outFilterMultimapNmax 20, --alignSJoverhangMin 8, --alignSJDBoverhangMin 1, --outFilterMismatchNmax 999, --alignIntronMin 20, --alignIntronMax 1000000 and --alignMatesGapMax 1000000. Additional parameter settings are--outFilterMismatchNoverReadLmax 0.04 and --outSAMattributes NH HI NM MD To visualize the mapped reads, bigwig files were generated using bamCoverage function in deepTools2 with the parameter setting as --normalizeUsingRPKM

### DNA-FISH probes

The bacterial artificial chromosomes (BACs) were obtained from the BACPAC Resource Center of Children’s Hospital Oakland Research Institute (Oakland, CA). DNA was isolated using BAC DNA miniprep Kit (Zymo Research). BAC probes were labeled using BioPrime Labeling Kit (ThermoFisher). Streptavidin, Alexa Fluor 488 conjugate (ThermoFisher, S-32354) was used to stain biotin-labeled BAC probes. Probes are described in Supplemental Table 2.

### 3-D DNA FISH/ immunocytochemistry and microscopy

3-D DNA FISH/ immunocytochemistry-labeling was performed as described previously (Vertii et al. 2019), except that DNA FISH-labeling was done after immunocytochemistry, and coverslips were not treated with RNA removal solution. F121-9 mESC were seeded on 0.1% gelatin-coated 22 × 22 mm coverslips (Corning, 2850-22), with total cell number 150-250 × 10^3^ cells/coverslip, and permeabilized and fixed the next day. Nucleoli were stained with anti-fibrillarin antibodies, except in the third biological replicates of the pPK999 and pPK1000 analyses anti-nucleophosmin antibodies were used instead.

Images were acquired using Zeiss LSM 700 laser scanning confocal microscope and PMT detector (63x 1.40 Oil DIC M27 Plan-Apochromat objective). DNA-FISH probes were counted through z-stacks manually and scored as “associated” if there was no gap between the probe and the nucleolar marker. Each probe was analyzed in at least three biological replicates, with at least 100 alleles scored in each replicate. Z stacks are represented as 2D maximum projections using Fiji software (Schindelin et al. 2012). Statistical analyses were done using GraphPad Prism software. p-values were calculated using arcsine values of the square roots of nucleolus-associated proportions.

### NAD identification and annotation

We used the same workflow for NAD-seq data analysis as described previously (Vertii et al. 2019), except that we removed 20 NAD peaks that are less than 50 kb long (totaling 0.74 MB). Because there are 624 peaks totaling 845 Mb in the F121-9 NAD-seq data, this represents 0.087% of the NAD nucleotides. We used version 1.6.1 of NADfinder for NAD identification in this manuscript.

Nucleotide-level overlap analyses of F121-9 NADs with cLADs, ciLADs (Peric-Hupkes et al. 2010), MEF NADs (Vertii et al. 2019), and H3K27me3-enriched domains (GSM2416833; (Cruz-Molina et al. 2017); GSM1621022; (Delbarre et al. 2017)) were performed using *GenomicRanges* (Lawrence et al. 2013) as described in detail in Vertii et al., 2019. These nucleotide-based overlap analyses in some cases generated small overlapped regions, such that single genes would end up with both Type I and Type II designations, or both MEF-specific and F121-9-specific designations. Because the biology of NADs is centered on large (∼1 MB-sized) domains, we removed regions <50 kb in length from overlap analyses of Type I and II NADs and from cell-type-specific NADs to avoid these confounding designations. GO enrichment analyses of conserved and cell type-specific NADs derived from the overlap analysis were performed using *ChIPpeakAnno*. mESC H3K27me3-enriched domains were identified based on H3K27me3 ChIP-seq data (GSM2416833; Cruz-Molina et al. 2017) using RSEG (v0.4.9) with 20 iterations for Baum training. MEF H3K27me3-enriched domains were obtained from GSM1621022 (Delbarre et al. 2017). FPKM values based on MEF RNA-seq data were obtained from GSM1621026 (Delbarre et al. 2017) and GSE90894 (Chronis et al. 2017). FPKM values from mES RNA-seq data were obtained from GSM1418813 (Lowe et al. 2015). Calculations of the statistical significance of pairwise comparisons were performed using Welch’s t-test in GraphPad Prism.

The *NADfinder* software is available at: https://urldefense.proofpoint.com/v2/url?u=https-3A_bioconductor.org_packages_release_bioc_vignettes_NADfinder_inst_doc_NADfinder.html&d=DwIFAw&c=WJBj9sUF1mbpVIAf3biu3CPHX4MeRjY_w4DerPlOmhQ&r=JqQ8_Clm34xp32rT3DzotqsofamUUUyNmo3M4_tlIEI&m=Lq6n57MH0XVDSsayaTs25TVTysYxezReg6cHQXKhVNk&s=BG-jkVe3qQRszk64lZLOGYCGqyYe-h9NoghI0r8I1bM&e=,

We calculated Jaccard indexes among NADs, cLAD/ciLAD (Peric-Hupkes et al. 2010), and F121-9 early/late replication timing (GSE95091 (Marchal et al. 2018)). The Jaccard index is the size of the intersect divided by the size of the union of two sets. The higher the Jaccard index, the higher the extent of the overlap.

Boxplots and comparisons of gene densities (genes/Mb) and gene expression distributions were performed using R For statistical comparisons, p-values were calculated using Welch’s t-test.

## Declarations

### Ethics approval and consent to participate

Not applicable.

### Consent for publication

Not applicable.

### Availability of data and materials

The datasets supporting the conclusions of this article are publically available at the 4D Nucleome Data Portal (https://data.4dnucleome.org/). The RNAseq data is at https://data.4dnucleome.org/experiment-set-replicates/4DNESDHILYLU/#raw-files. The DNAseq data for the total genomic samples is at https://data.4dnucleome.org/experiment-set-replicates/4DNESUJZ5FL2/, and the DNAseq data for the nucleolar samples is at https://data.4dnucleome.org/experiment-set-replicates/4DNESXE9K9DB/.

### Competing interests

The authors declare that they have no competing interests.

### Funding

Research reported in this publication was supported by the National Institutes of Health (National Institute on Drug Abuse, U01 DA040588 to P.D.K.) as part of the 4D Nucleome Consortium.

### Authors contributions

AB performed all the wet lab experiments, and analyzed and interpreted the data. AY and JY performed bioinformatic analyses. LJZ directed and conducted the bioinformatics analyses, and PDK directed and analyzed the wet lab experimentation. AB, LJZ and PDK wrote the manuscript. All authors read and approved the final manuscript.

## Acknowledgements

We thank Takayo Sasaki and David Gilbert (Florida State University) for the kind gift of F121-9 cells, and Anastassiia Vertii for guidance with the nucleolar preparation and DNA-FISH experiments.

## Supplemental Tables

Supplemental Table 1: Average RNA-seq FPKM values from two biological replicate RNA-seq samples, made from the same preparations of F121-9 cells used for the nucleolar purifications.

Supplemental Table 2: mm10 genomic coordinates, laboratory BAC probe names, systematic BACPAC names, FISH and NADfinder results for DNA-FISH probes.

Supplemental Table 3: GO-derived Molecular Functions terms of conserved NADs, with q-values (termed “BH adjusted p-value” in the table) below 0.05.

Supplemental Table 4: GO-derived Molecular Functions terms of F121-9-specific NADs, with q-values below 0.05.

Supplemental Table 5: GO-derived Biological Processes terms of MEF-specific NADs, with q-values below 0.05.

Supplemental Table 6: mESC H3K27me3-enriched domains identified based on H3K27me3 ChIP-seq data (GSM2416833; Cruz-Molina et al. 2017) using RSEG software. The first three columns show for each H3K27me3-enriched domains the chromosome name, start and end nucleotides. The 4th column (Average Count) gives the average read count in the domain. The 5th column (Domain Score) is the sum of posterior scores of all bins within this domain; it measures both the quality and size of the domain.

## Notes

#### Summary of Updates

We have added declarations regarding ethics approval, competing interests, funding, authors contributions, and acknowledgments.

https://data.4dnucleome.org/experiment-set-replicates/4DNESDHILYLU/#raw-files

https://data.4dnucleome.org/experiment-set-replicates/4DNESUJZ5FL2/

https://data.4dnucleome.org/experiment-set-replicates/4DNESXE9K9DB/

